# Memory Inception through Gaze-Contingent Message Exposure: Using Virtual Reality to Study Media Influence

**DOI:** 10.1101/2025.01.08.631803

**Authors:** Hee Jung Cho, Sue Lim, Miriya Saenz, Ralf Schmälzle

## Abstract

The messages we encounter in our environment can shape our knowledge about the world. However, much research on media-driven influence via messages focuses on population-level effects and aggregate exposure statistics, obscuring how individual and self-determined behaviors affect message intake, processing, and effects. To address this gap, we use virtual reality (VR) to create a controlled messaging environment. Participants navigate a simulated urban street lined with billboard messages while their visual attention is tracked via eye-tracking. We introduce an inception-style manipulation: overlooked billboards are strategically reintroduced, creating additional exposure opportunities. Our results demonstrate that this subtle manipulation – unnoticed by participants – boosts message retention. This study bridges communication theory and psychology, elucidating the blurred line between voluntary and involuntary information intake in the digital age. It also highlights a potential vulnerability in the future metaverse media ecosystem, where undetected information manipulations can influence individual and collective attention and memories.

## Main

What if one could slip an idea into someone’s head, like in the movie Inception where agents infiltrate and manipulate a person’s memory system? While the film leaves its technical mechanisms mysterious, the concept of similar cognitive manipulations has long fascinated scholars across disciplines. Researchers have explored this through biotechnological studies of memory transfer (Rosenblatt et al., 1966; Domjan, 2014), investigations of media propaganda and persuasive techniques (Lasswell, 1927; Skinner, 1958; McCombs and Shaw, 1972; Herman and Chomsky, 1988; Turow, 2022), or even studies of beliefs about “influencing machines” (Tausk, 1919; Gladstone and Neufeld, 2011). These studies underscore a central question: How and to what degree can external forces like media messages and technological manipulations shape human memory?

While the concept of remote-controlled influencing machines remains science fiction, real-world media can subtly shape our minds by delivering messages that enter through our senses, ultimately influencing our thoughts and behaviors (Zillmann and Bryant, 1985; Schmälzle and Huskey, 2023). Over a century of research demonstrates that we form memories based on the information we encounter in our environment, whether in the form of messages like billboards, internet banners, TV ads, or simply word lists in laboratory experiments (Ebbinghaus, 1885; Gallistel and King, 2011). This aligns with the concept of exposure in mass communication research, which represents the initial step in the process from message transmission to media effects (Neijens et al., 2024). High-profile events like the Super Bowl exemplify the ability of media to reach a mass audience and expose them to advertisements that leave tracks in collective memory (Hartmann and Klapper, 2018). However, the same principles apply to other media formats and a wide range of communication objectives – from increasing brand awareness to advocating for specific causes, all of which ultimately aim to influence the minds of individual message recipients (Valkenburg et al., 2016).

A persistent challenge in media exposure research is the reliance on aggregate metrics like TV viewership size, billboard traffic, or online statistics, which are prone to measurement error and manipulation (Nelson and Webster, 2016). These metrics mistakenly assume that message availability equates to message intake, overlooking the fact that widespread broadcasting does not guarantee individual attention or processing. Instead of focusing on the mere availability of messages, valid measures of exposure should track message reception. For visual messages, eye-tracking thus offers a more accurate assessment of exposure as it happens (Kingstone et al., 2003). By adopting such an objective and micro-level perspective, we can gain a deeper understanding of how individuals engage with and are influenced by the messages they encounter (Zillmann and Bryant, 1985).

Challenges in studying message exposure in ecological settings have hindered the integration of an information-processing perspective, which is standard in lab-based memory experiments, to the complexities of media research. To overcome this gap, recent work has begun to use virtual reality (VR) to study how people encounter and encode messages under conditions that resemble real-life messaging environments (Bonneterre et al., 2024; Clay et al., 2019; Schmälzle et al., 2023). These studies examined how participants self-select messages based on their attention patterns as they navigate virtual worlds, such as cities or highways. Findings show that overt visual attention, like fixating on billboards, gates message intake and influences memory retention. Distractions, by reducing fixations, weaken memory (Cho et al., 2024; Lim et al., 2024). This highlights how attention transforms exposure opportunities into actual exposure, acting as a gateway and bottleneck for information to enter the mind, which forms the basis for memory (Broadbent, 1958; Neisser, 1964; Potter, 2008; Sherman and Turk-Browne, 2024).

### Potential to create Inception-Style Media Exposure Paradigms

If attention serves as a gatekeeper under everyday exposure conditions, what happens when this gate is manipulated? The studies discussed above suggest that messages that are at least briefly fixated on are far more likely to be remembered than those that are ignored. This raises the question: how can we influence attention in a way that tilts the scales in favor of or against it? Cognitive psychology suggests several avenues, such as increasing physical saliency (e.g., Kümmerer and Bethge, 2023), adding affective content to the message (Schupp et al., 2004), or leveraging receiver-sided variables, such as motivational or task relevance (Todd and Manaligod, 2018). Each of these factors should affect the attentional gate, and modern advertising techniques clearly seek to exploit such ideas (Armstrong, 2010). Accordingly, the prior VR-eye-tracking studies also sought to influence this exposureattention-gate in different ways – from making messages more salient, placing them more conspicuously, or by exhausting recipients’ attentional capacities (Bonneterre et al., 2024; Cho et al., 2024; Jeon et al., 2024; Lim et al., 2024). Despite these efforts, however, attention remains a somewhat unpredictable, fluctuating process (Esterman and Rothlein, 2019), with the decision to engage with a message still largely in the hands of the individual. As John Wanamaker, a pioneer of modern advertising, famously remarked: “Half the money I spend on advertising is wasted; the trouble is I don’t know which half.”

The challenge, then, becomes: could we use a side-gate into the nexus between exposure and attention? Can we reduce the uncertainty of self-deployed and wandering attention to ensure that a message receives the attention necessary for memory encoding? This is where the “inception paradigm” comes into play, which refers to a system that increases the likelihood of actual exposure through dynamic adjustments to the messaging/media environment. The core of this paradigm consists of a mechanism that allows messages to be reintroduced until they are attentively viewed, a bit like incepting an idea into the mind.

To explain, consider a driver navigating a virtual city, passing various billboards along the way. The key is to track whether the driver looks at a given message. If they do, then we know that the message has received at least a minimal dose of attention and thus should be more likely to be remembered. However, if they do not, this is where the innovation lies: Using VR-integrated eye-tracking, we can detect a failed exposure—a message that is passed without being looked at. Rather than allowing this message to be disregarded as a failed exposure, it can be reintroduced further down the road. Then, by repeating a message that was initially ignored, we have a chance to transform a failed exposure opportunity into a successful actual exposure, which should heighten the probability of encoding the message into memory.

Critically, such gaze-contingent message exposure is no longer a concept of science fiction: Because the urban environment, including the road and the billboards, are all digitally represented in VR and eye-tracking data is constantly monitored, this can be accomplished algorithmically using current technology (see Figure 1 and Materials and Methods).

**Figure 1:**
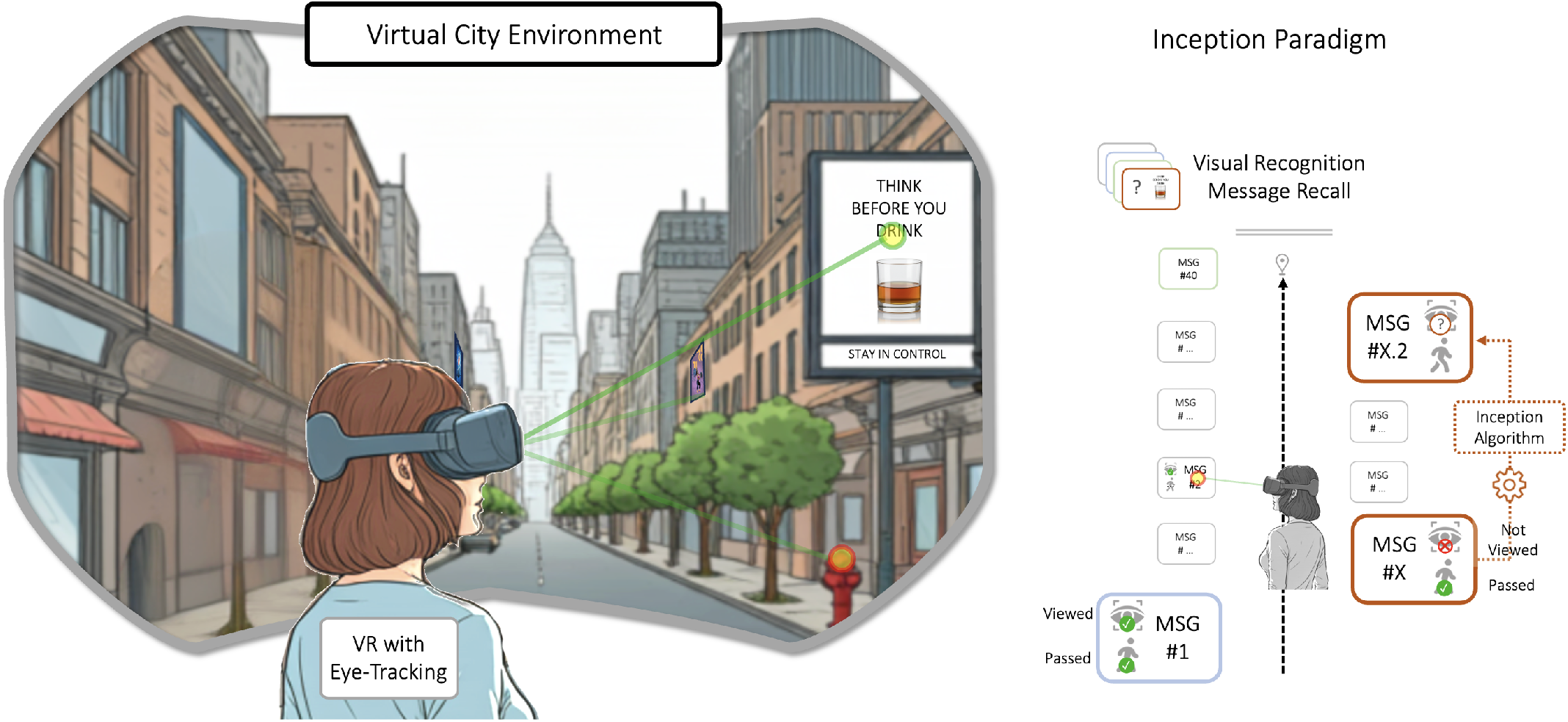
Study Design. A virtual city serves as a visual communication environment to study exposure to and reception of real-life messages. Via VR-integrated eyetracking, we objectively assessed whether participants looked at each message as they passed it. Messages that were passed but not looked at could be algorithmically re-shown, thereby creating another opportunity for exposure to selectively target memory for those messages. Once participants reach their destination, their retention of messages is measured, allowing to reveal the effects of self-deployed attention and incidental memory and the success of the inception attempt.

To test these ideas, we expanded our previously validated system for studying message exposure in an urban environment (Schmälzle et al., 2023; Lim et al., 2024). This system was adapted to make algorithmic decisions, re-showing messages to participants who had initially missed them. The critical experimental intervention consisted of taking a subset of messages that were passed but not viewed and presenting them again. We predicted that this should boost the likelihood of those messages to be remembered.

A crucial follow-up question was whether participants would even notice this modification. In essence, the proposed inception paradigm extends principles of message targeting by targeting the attention of recipients, although it resembles more a “catching” than a direct targeting of attention. This may make the technique less noticeable and perhaps more effective than conventional targeting, which is frequently noticed, perceived as creepy, and not all too effective.

Studying whether this attempt to influence memory is detected is also relevant as we move towards metaverse-like communication environments, where undetected information alterations could potentially be applied on a large scale, and perhaps even more subtly than what we know from the internet (e.g., Kramer et al., 2014; Matz et al., 2017). Lastly, we sought to replicate prior findings regarding the impact of distraction on attention patterns and the link from attention to message retention.

## Results

Participants (*m*_age_ = 19.8; *sd*_age_ = 1.3; 24 female) completed the VR-based drive through our experimental city, which contained forty professionally designed billboard messages along the way. The messages featured a variety of billboard-typical topics, including commercials (e.g. burger restaurants, hotels, consumer items and services) as well as various health-related public service messages (e.g. texting and driving, risky alcohol use, smoking cessation), and were randomized to positions throughout the city. Twenty participants were assigned to the free viewing condition and instructed to simply drive down the road. Twenty other participants were assigned to a visual distraction condition and instructed to count the number of trash bins they could spot in the city. Upon exiting the VR simulation, we conducted an interview as well as an unannounced message recall test, followed by a survey probing recognition memory and participants experiences in VR (see Figure 2a).

**Figure 2:**
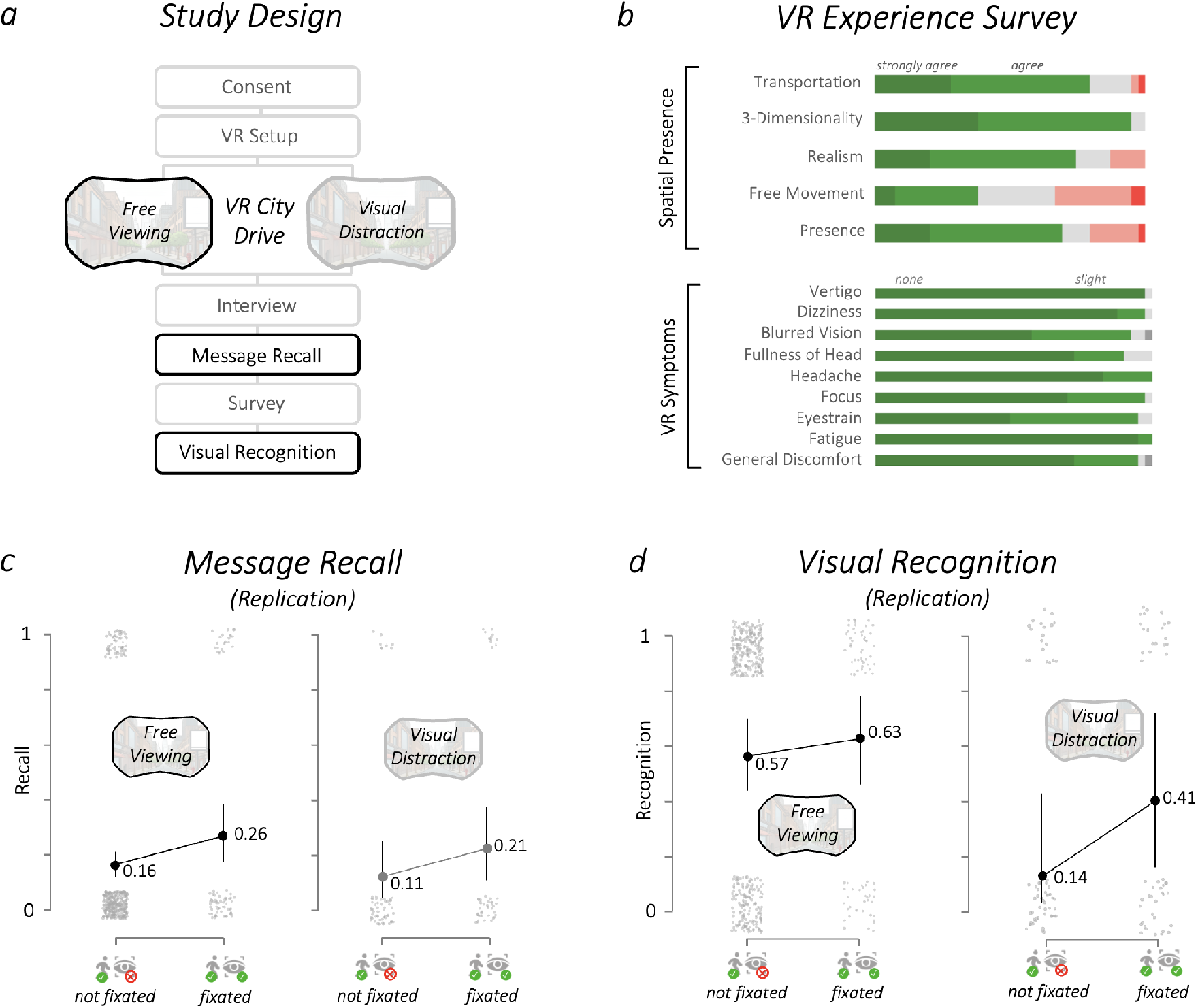
Study Design, VR Survey Results, and Replication of the Attention-Retention-Gating Effect. a) Study timeline and conditions. b) Survey results confirm that participants felt present in the VR city environment and experienced little to no symptoms due to VR. c) As in prior studies, participants were far more likely to recall fixated messages, and this was seen in both conditions (free viewing, with more fixations) as well as the trash-can-counting distraction task. d) Same results as in panel c, but for recognition memory.

### Assessing immersive experience during the VR city drive

In immediate post-VR interviews, participants commonly described the experience as smooth and immersive, often highlighting its interesting and pleasant nature. Survey results supported these evaluations, showing a high level of spatial presence (*mean*_spatial-presence_ = 3.78; *sd* = 0.58; Hartmann et al., 2016), which is significantly above the scale midpoint (range 1-5; *t*(39) = 8.29, *p* < .001). Participants also reported very little to no physical discomfort due to VR (*mean*_VR-symptoms_ = 1.3; *sd* = 0.32; Kim et al., 2018). This is significantly below the scale midpoint (range 1-4; *t*(39) = −23.9, *p* < .001), and all individual symptoms such as general discomfort, fatigue, and dizziness were rare, with average symptom levels ranging from none to slight, and all significantly below midpoint (*p* = .001; see Figure 2b). This combination of high spatial presence and low discomfort suggests that participants felt comfortable and explored their surroundings naturally, making it a suitable model for studying human behavior in both real-life as well as metaverse-style communication environments.

### Participants are not aware of the manipulation

To evaluate whether participants noticed the experimental manipulation (re-showing of billboards based on participants’ fixation patterns during the drive), we asked them directly whether they observed anything unusual. Thirty-nine out of 40 participants (97.5%) did not mention anything. One participant mentioned spotting a flicker and a change in a sign toward the end of the drive, but even this participant failed to recognize that the same sign had been re-shown, or that multiple other billboards had been altered. Thus, we conclude that the manipulation was not detected.

### Replicating the gating effect between attention and message retention

Having established that participants experienced the virtual city drive as realistic and were not aware of the experimental manipulation, we next turned to the effect of attention on memory recall. First, to replicate the finding that overt visual attention on billboards would impact retention, we examined recall and recognition memory for billboards that were fixated during the first passing and compared them to billboards that were not fixated during passing and then never shown again.

As expected, and shown in Figure 2, we find a strong effect of attention: fixating a billboard drastically boosts the likelihood that it will later be recalled from near-zero recall rates for billboards that were passed but not fixated to about 20% recall for those that received at least a minimum dose of attention. Statistical assessment of the results via generalized linear mixed models revealed that messages that were passed and attended to were far more likely to be recalled than those that were passed but not fixated (χ^2^_*Attention Status*_ = 5.654, *p* = .017). There was no significant interaction with viewing condition, though nominally the recall was higher in the free viewing condition, which also resulted in more fixations. For visual recognition memory, a very similar pattern emerged: Memory for messages that were passed and attended to was significantly higher compared to those that were passed but not fixated (χ^2^_*Attention Status*_ = 4.416, *p* = .036) raising recognition rates from about 15-20% up to 50-70%. For recognition memory, there was a significant main effect of condition (χ^2^_*Viewing Condition*_ = 5.897, *p* = .015); those in the free viewing condition recognized more billboards than those in the distraction condition. There was no significant interaction (χ^2^ = 1.402, n.s.).

### Message inception effect

The most critical comparison is between re-shown billboards that were then fixated vs. those that were not. These billboards were re-shown because they had not been looked at during the first passing. Consistent with our hypothesis that a dose of overt attention to a billboard would markedly alter its fate in memory, we find that billboards that were fixated during a re-showing achieved recall rates of about 60% (during free viewing) or 30% (trash-bin-counting, see Figure 3a). The GLMM revealed a main effect of message viewing status (χ^2^_*Attention Status*_ = 70.409, *p* < .001) on message recall. There was no significant main effect of condition, although the free viewing condition nominally garnered more fixations and was associated with higher memory. There was also no significant interaction between viewing status and condition.

**Figure 3:**
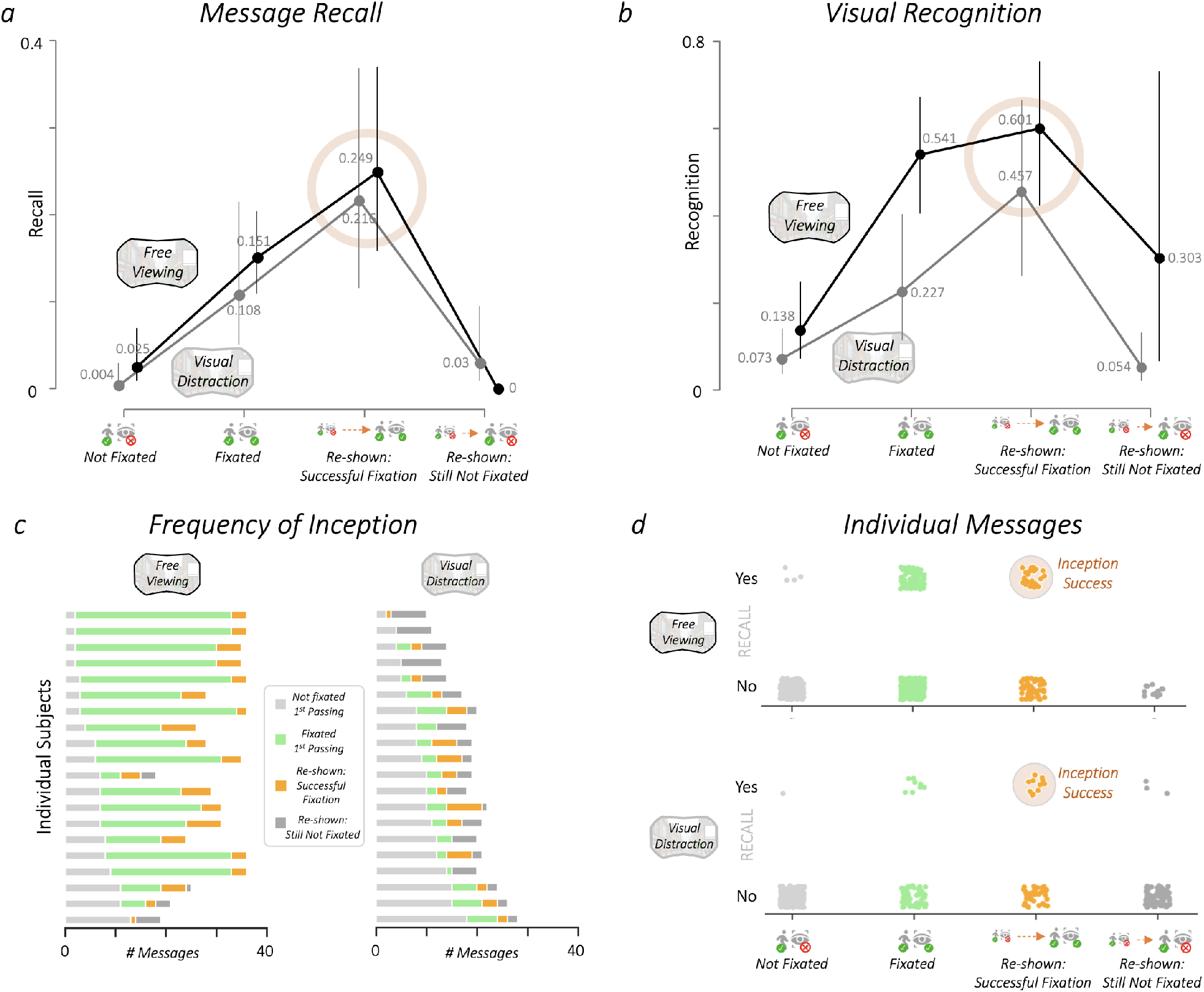
Inception Effect. a) A subset of passed messages that were not fixated were shown again in the hope to attract eye gaze. If successful, i.e. if the messages were fixated, then they are substantially more likely to be recalled. This effect is consistent across the free viewing and visual distraction conditions. b) Same as in A, but with message recognition as outcome. c) Due to the self-determined nature of participants’ viewing behavior, inception attempts could only be triggered if a participant passed a billboard but did not fixate it. Shown are the distributions of viewing categories for all participants and conditions. d) Swarmplots of all individual messages, organized by viewing status and memory outcome (recall). As can be seen, messages that were fixated, are more likely to be remembered. Critically, the re-shown messages garnered many additional fixations, which in turn lead to these messages being more likely to be recalled.

Post-hoc testing confirmed that messages that were re-shown and fixated were significantly more likely to be recalled compared to the baseline messages, i.e. those that were passed and not fixated, and then not selected for inception to serve as baseline (χ^2^ = 58.469, *p* < .001). Re-shown and then fixated messages were also more likely to be recalled compared to re-shown messages that still failed to attract attention (χ^2^ = 16.871, *p* < .001).

A similar pattern of results emerged for recognition memory (Figure 3b): Successfully attracting attention due to reshowing a previously unfixated billboard boosted recognition memory from ca. 10-20% for unfixated billboards to about 3060% for the inception condition. This gain was higher in the free viewing condition, but still substantial even in the trashbin-counting condition, which had generally less fixations. Statistical analysis revealed again a highly significant main effect of attention status (χ^2^_*Attention Status*_= 78.438; *p* < .001), with a significant main effect of viewing condition (χ^2^_*Viewing Condition*_ = 7.163; *p* < .001), but no significant interaction effect. Again, subsequent comparisons confirmed better recognition memory for the successfully reshown messages compared to both the baseline messages (passed, unfixated, and not re-shown, χ^2^ = 63.089; *p* < .001) as well as unsuccessfully re-shown messages (i.e. re-shown, but unfixated; χ^2^ = 22.593; *p* < .001).

The results demonstrate that the inception manipulation worked at the group level, and consistently for both the free viewing as well as the distraction (trash-bin-counting) condition. This underscores the viability and potential of the approach. However, an important open question concerns the relative success rate and impact of inception attempts. Answering this question is not completely straightforward though, because whether and when an inception attempt is triggered depends on each participant’s viewing behavior, which varies considerably. For instance, if a participant looked at all billboards, then there would be no inception at all. Moreover, the viewing task (free viewing vs. distraction via trash-counting) also impacts viewing behavior, and thus the frequency of inceptions.

Therefore, we closely examined participants’ viewing behavior (Figure 3c) as well as results for individual messages (Figure 3d). In this study, participants in the free viewing condition fixated on average 19.35 billboards during first passing, about half of the 40 billboards. In the distraction condition this number was far lower, with only 3.05 billboards reaching fixation status (about 10%). Accordingly, the distraction-condition triggered far more inception attempts, but these attempts should also be less likely to find success because participants kept searching for targets. Indeed, in the distraction condition, an average of 6.35 inception attempts was made, and those had a 37% chance of finding success (i.e. leading to a fixation on a billboard that was previously passed without fixating). In the free viewing condition, on the other hand, there were an average of 4.65 inception attempts, but these had a far higher success rate of 87%.

Relating these metrics back to the original argument that many message dissemination efforts remain futile, we can see that for the free-viewing condition, we were able to boost effective exposure from about 50% to about 60%. Also, it should be noted that we opted to not re-show all unfixated messages but only a subset thereof and use the rest as a baseline for memory without fixations. If this inception-system were actually implemented, one would drop this aspect and instead re-show all unfixated messages to push effective exposure rates even further. According to recent data, global outdoor advertisers spent over $40 billion on messaging. Against this backdrop, achieving an increase in efficiency of more than 10% is clearly a major factor. Moreover, these calculations apply only to outdoor advertising, not even considering the colossal digital advertising market to which they equally apply.

## Discussion

Our findings reveal that the inception manipulation effectively transformed many failed exposure attempts into successful message encounters. This increased both retention of message content as well as familiarity. Participants did not notice how the inception algorithm effectively steered the information they were served, which in turn embedded messages into their memory that otherwise would not have formed. Successfully incepted billboard messages achieved recall rates of about 20% and recognition rates of up to 60%, compared to far lower rates for overlooked messages.

These results contribute to the ongoing debate about measuring exposure in mass communication and media research, with a particular emphasis on the blurry boundaries between voluntary and involuntary information intake in the attention economy. In today’s world, we are inundated with thousands of messages per day – from billboard ads during our commute, to social media posts, to TV messages. Interestingly, while we often recognize the enormous influence media seem to have on others, like when children imitate role models or adults parroting message frames (Bandura, 2009; Entman, 1993), we tend to underestimate its impact on ourselves (Perloff, 1999). Although we did not specifically study so-called third-person effects in this investigation, we believe that the self-determined nature by which we encounter the information may at least in part contribute to them.

Critically, participants in this study remained unaware of our inception manipulation and formed no suspicion that part of their messaging was dynamically targeted. Their perception was that they were free to visually explore their surroundings, and they indeed could look wherever they wanted; but behind the scenes our inception algorithm pulled the strings as to which messages would appear in front of their eyes, which shapes what they remembered. Therefore, our inception-manipulation does not amount to a deterministic “steering” of attention towards every message. Rather, it operates by modifying the environment to generate conditions that favor exposure to specific messages and their subsequent retention. Despite this reservation, this gets as close as possible to causality without creating a kind of “forced exposure” experiment as is common in laboratory research on intentional memory (Ebbinghaus, 1885), but which would no longer resemble the kinds of visual media environments we encounter and from which we form incidental memories.

Whether we notice how we are influenced by messages or not, there is no question that the myriads of incidental message exposures can shape our thoughts, highlighting the power of media to impact individuals and society at large. Because message effects are contingent on prior exposure, exposure has been dubbed the foundation of all media effects (McGuire, 1968; Slater, 2004). The innovation of our approach is that it turns a variable that is measured – often incompletely and only at an aggregate level – into one that can be precisely isolated and manipulated. Objective measurement and control have always been major challenges in the social and behavioral sciences. In this context, technology-mediated manipulations and transformations like our VR-based approach to study message reception are creating powerful tools for social scientists (Blascovich et al., 2002; Miller et al., 2019). This is also underscored by similar work demonstrating how eye-gaze or facial expressions can be technologically manipulated to bias consequential social decisions (Arias-Sarah et al., 2024; Pärnamets et al., 2015).

We note that exposure as it is studied in communication and media research differs from the mere exposure effect in psychology (Zajonc, 1968), though several connections are worth noting. In brief, the mere exposure effect refers to the phenomenon where people tend to develop a preference reaction because they are made more familiar with stimuli. However, looking into the literature on the mere exposure effect (Bornstein and Craver-Lemley, 2022), it is clear that what was manipulated is largely the number of showings of the stimulus (e.g. people appearing more frequently in a class over a semester, or foreign signs being flashed more often, etc.). This again points to the distinction between making a stimulus or message available in the information environment (exposure opportunities) and measuring the actual intake by the human cognitive system (actual message encounter). Even so, work on the mere exposure effect is relevant insofar as it is often used as a basis for brand awareness and other core concepts in consumer psychology and media effects research (Sutherland, 2020).

Moreover, our results can be interpreted in light classical mere exposure research because the messages in the inception condition, which were necessarily presented multiple times, were in fact significantly more likely to be recalled and recognized than messages that were just passed and looked at once (see Figure 3). However, we also find that if the re-shown messages were not looked at, their memory advantage vanishes. This again underscores the role of attention in converting exposure opportunities into actual message reception (e.g. Broadbent, 1956). In contrast to mere exposure research, we did not examine whether the messages people viewed and remembered were more liked on an attitudinal level, but we believe our study can also help trace the pathway by which mere exposure effects occur.

Finally, considering practical and societal implications, these are obvious and significant. The ability to turn exposure opportunities into actual exposure via VR-based inception speaks directly to the advertising-based business model of media giants like Meta, Google, Tencent, and others. Especially given the predicable growth of VR-mediated communication environments, like the Metaverse, and the advances in commodified VR headsets that include eye-tracking, this work also has implications for user privacy and data protection (Farahany, 2023; Turow, 2017).

We acknowledge certain limitations. First, we focus on shortterm memory, leaving open questions about how lasting the effects are. Given the literature repeated exposure to messages (Skurka and Keating, 2024) and the influence of repetition on memory in list-learning studies (Ebbinghaus, 1885), we believe it is justified to focus on short-term memory and extrapolate. However, we note that if the inception attempts were repeated, people would also start becoming more aware of them, much like we get uncanny feelings when targeted ads seem to follow us across platforms. Second, we also acknowledge that other eye-tracking metrics, such as rapid saccades and short glances, can offer more fine-grained insights, as even sort glances could still be enough to form gist-based memory (Lleras et al., 2022). Third, given that our goal was to demonstrate the feasibility of memory inception, we only focus on message memory as an outcome. Thus, it remains open whether observed effects extend to other outcomes, most notably attitudinal and behavioral ones as in mere exposure studies.

Going forward, research should examine the scope of these inception-style influences, particularly whether it is possible to impact more complex ideas or frames (e.g. Coronel et al., 2023); another avenue is to focus on deeper and more longer-term outcomes. Efforts could also be expanded to more complex and dynamic environments by incorporating a broader range of user states and actions such as approach or avoidance behaviors. For instance, the display of ads could be made contingent on motivational state variables (e.g. showing more food ads to hungry subjects). The way the internet evolved has clearly shown that digital traces enable inferences about users’ interests and personality, which are then used for messaging (e.g. Matz et al., 2017). Therefore, it becomes a pressing research priority to examine the impact of these manipulations on consumers and their ability to detect and protect against them.

### Conclusion

We developed a novel method – a gaze-contingent algorithm for re-display of unattended messages in VR – that allows studying, measuring, and experimentally manipulating exposure in realistic and new media environments. Our findings reinforce the connection between exposure and memory and open up possibilities for precise message delivery, with significant implications for society.

## Methods

### Participants

We assigned twenty participants to a free viewing condition and another twenty to a visual distraction condition. Thus, there were 40 participants in total (*m*_age_ = 19.8; *sd*_age_ = 1.3; 24 female). Three additional participants were recruited but failed to complete the study due to technical reasons and were immediately replaced. All participants received course credit and provided written informed consent to the study, which was approved by the local Institutional Review Board.

### Materials and Equipment

#### Stimuli. Billboard Messages

We used 40 billboard messages as experimental stimuli. Based on previous work (Lim et al., 2024), we adapted 20 distinct health and risk-related topics (e.g. smoking, vaccination, texting and driving) and 20 commercial topics (e.g. hotels, restaurants, lawyers). The messages were professionally designed to feature a mix of visual and textual components typical for the genre of outdoor advertisements/PSAs (see Figure 1 and Supplementary Materials for examples). Once we generated both the slogans and images (via ChatGPT and Midjourney services), we used an 800 × 800 Canva template to format them to reflect a realistic outdoor billboard.

#### Virtual Environment

We used Vizard’s Sightlab VR Pro Software (Worldviz.com) to create and display the VR city driving experience and measure gaze behavior as participants navigated the true-to-scale visual environment. Specifically, the “Classic City: Mobile” 3D model was downloaded from the Unity Asset Store (see Lim et al., 2024). This environment features common city structures, such as streets, buildings, and typical city-like accessories like trash bins, lamps, trees. To reduce potential distractions, we made modifications to the city and allocated 40 spots for advertisements on buildings and in parks along the city streets. These placements were distributed fairly evenly on both sides of the street, with slight variations in height, distance from the road, and angle to enhance the realism.

#### VR Head Mounted Display (HMD) with Integrated Eye-tracker and Navigation

The HP Reverb G2 Omnicept served as VR head-mounted device (HMD). This HMD features a high-precision integrated Tobii eye-tracker. Participants used the VR controller to drive straight down the city street, with the right trigger button serving as gas pedal and the left trigger as the brake. They could accelerate up to a speed of about 20 mph, leading to a smooth cruise down the road. To decrease the confusion and ensure that participants simply drove down the street with the 40 billboards, there was no need for the participants to steer.

#### Procedures

The experiment began with participants arriving at the laboratory and completing the informed consent process. After signing the necessary documents, subjects underwent a quick vision assessment before donning the virtual reality headset. The research team then calibrated the integrated eye-tracking system and provided instructions on using the VR controller. Participants were informed they would be exploring a virtual cityscape that was temporarily closed for maintenance.

The cohort was divided into two groups: one tasked with tallying trash cans (Trash can counting: n = 20), and another instructed to simply drive without further instruction (Free viewing: n = 20). The virtual driving experience lasted approximately 10 minutes. Upon completion, researchers conducted brief interviews to gather impressions of the simulated drive and assess participants’ recall of advertisements encountered. This was followed by a Qualtrics survey covering demographics, presence perception, simulator sickness, and message recognition. The session concluded with a thorough debriefing of all participants.

#### Detection of Eye-Gaze towards Billboards and Experimental Inception Algorithm

In brief, as participants drove down the virtual street, the VR-integrated eye-tracker tracked their viewing behavior, and the VR software recorded their relative position on the road. Thus, we can determine whether a driver who passed a billboard fixated it or not. If the driver passed a billboard and had fixated it, this message was added to a list of messages that were fixated upon the first passing. However, if the passed billboard had not been fixated, a random choice was made between two options: First, the passed billboard could either be added to the list of “forgone” billboards, which served as a baseline control (assuming, based on prior work, that billboards that were not looked at, would not be remembered). The second-choice option – and our main experimental manipulation – was to set the billboard up as one to be incepted.

Programmatically, this “inception attempt” works as follows: Assume a driver passed by the first billboard (Billboard1 at Position1) but didn’t fixate it. The algorithm chooses between using the billboard as control or scheduling it for inception. Assuming the “to-be-incepted” option is chosen, then the program will swap a billboard positioned down the road, specifically at position X+5 (i.e. the 6th billboard in this example). Given that the 5th billboard couldn’t be seen from the vantage point of position1, the driver will not know that this has happened. As an analogy, this is as if people were conspiring behind the scenes to telegraph a person located at position 6 to swap out the original billboard6 with a copy of the billboard that was originally shown at position1.

In essence, this inception manipulation creates a second chance for the driver to look at billboard1 – now shown on position6. However, if the driver again passes billboard1 at position6, yet still doesn’t incidentally view it, then the program will create another option for inception – i.e. inception attempt 2 at the position x+5, now position 11. This is repeated up to three times, at which point the inception attempt is abandoned if the billboard has still not attracted a fixation.

As a result of this manipulation, there are four categories of viewing status: First, the billboards that were passed but not viewed and that were added to the forgone list to measure baseline memory. Second, the billboards that were fixated upon first passing. Third, the billboards that were re-shown (inception attempt) and then actually looked at. Fourth, billboards that were re-shown but still not looked at. In sum, this inception procedure experimentally creates additional opportunities for exposure, allowing us to test if this manipulation increases the number of fixations to billboards and subsequently memory.

#### Measures

##### Gaze behavior

We used Python in Vizard to track fixations and gaze duration for each billboard, with a fixation threshold of 0.25 seconds. Moreover, because the eye-tracker sampled participants’ gaze continuously, we also kept track of sub-fixation ‘glances’ at individual billboards for later inspection. Thus, the main variable we’re interested in is whether a participant devoted at least minimal visual attention to a given billboard (i.e. fixated it).

##### Message Recall

At the end of the virtual drive, after removing the headset and answering brief interview questions about their experience, we conducted an unannounced test of participants incidental memory. Specifically, we asked them to freely recall as many billboards as possible.

##### Visual Recognition

In addition to the free recall measure, we also assessed participants’ recognition memory. To this end, a final survey was administered via Qualtrics software on an iPad, showing participants images of all experimental billboards as well as X distractors, and asking them to indicate whether they recognized the billboards as having been part of the cityscape they just navigated (yes/no).

#### Data Analysis

The data analyses were conducted using Jupyter Notebook for data preparation and JASP software (JASP Team, 2024) for statistical analysis. Memory outcomes including recall and recognition, were binary-coded (recalled/recognized = 1, not recalled/recognized = 0). For the recognition test, we also included 4 dummy distractor billboards (i.e. billboards that actually did never appear in the city) in order to catch guessing strategies and gauge participants’ tendency for false recognition or However, we observed that participants did not just pretend to recognize all billboards, but rather responded variably to the recognition questions; also, on average only one in twenty indicated having seen a dummy distractor (due to false memory/guessing), and no participant falsely reported having seen more than one dummy billboard. This demonstrates that participants did not engage in guessing but reported actual memories.

To test whether our results replicated prior findings, we first conducted generalized linear mixed model analyses, examining effects of driving conditions (free-viewing vs. visual distraction via trash-bin-counting) and attention (i.e. messages that were passed but not fixated vs. messages that were passed and overtly looked at) on memory. These models were computed using either free message recall or visual recognition as the outcome. Then, in the main analysis, we included also the incepted messages, leading to a variable viewing status comprising four levels (passed and not looked at, passed and looked at, re-shown and fixated, re-shown and still not fixated) as well as the factor viewing condition. Again, both memory outcomes (recall and recognition) served as dependent variables.

## Acknowledgements

We thank Sado Rabaudi (WorldViz, Inc.) for helping with the Sightlab system.

## Data availability

Anonymized data is available via GitHub at https://github.com/nomcomm/vr_billboard_p.

## Code availability

Code is available via GitHub at https://github.com/nomcomm/vr_billboard_p.

## References

Arias-Sarah, P., Bedoya, D., Daube, C., Aucouturier, J.J., Hall, L., Johansson, P., 2024. Aligning the smiles of dating dyads causally increases attraction. Proceedings of the National Academy of Sciences 121, e2400369121. doi:10.1073/pnas.2400369121.

Armstrong, J., 2010. Persuasive advertising: Evidence-based principles. Springer.

Bandura, A., 2009. Social cognitive theory of mass communication, in: Media effects. Routledge, pp. 110–140.

Blascovich, J., Loomis, J., Beall, A.C., Swinth, K.R., Hoyt, C.L., Bailenson, J.N., 2002. Immersive virtual environment technology as a methodological tool for social psychology. Psychological Inquiry 13, 103–124.

Bonneterre, S., Zerhouni, O., Boffo, M., 2024. The influence of billboard-based tobacco prevention posters on memorization, attitudes, and craving: Immersive virtual reality study. Journal of Medical Internet Research 26, e49344.

Bornstein, R.F., Craver-Lemley, C., 2022. Mere exposure effect, in: Cognitive illusions, pp. 241–258.

Broadbent, D., 1958. Perception and Communication. Pergamon Publishers.

Broadbent, D.E., 1956. Successive responses to simultaneous stimuli. Quarterly Journal of Experimental Psychology 8, 145–152.

Cho, H.J., Lim, S., Turner, M.M., Bente, G., Schmalzle, R., 2024. Eyes on vr: Unpacking the causal chain between exposure, reception, and retention for emotional billboard messages. bioRxiv doi:10.1101/2024.07.19.604208.

Clay, V., König, P., Koenig, S., 2019. Eye tracking in virtual reality. Journal of Eye Movement Research 12.

Coronel, J.C., Ott, J.M., Hubner, A., Sweitzer, M.D., Lerner, S., 2023. How are competitive framing environments transformed by person-to-person communication? an integrated social transmission, content analysis, and eye movement monitoring approach. Communication Research 50, 3–29. doi:10.1177/00936502209035.

Domjan, M., 2014. The Principles of Learning and Behavior. Nelson Education.

Ebbinghaus, H., 1885. Über das Gedächtnis: Untersuchungen zur experimentellen Psychologie. Duncker Humblot.

Entman, R.M., 1993. Framing: Toward clarification of a fractured paradigm. Journal of Communication 43, 51–58.

Esterman, M., Rothlein, D., 2019. Models of sustained attention. Current Opinion in Psychology 29, 174–180. doi:10.1016/j.copsyc.2019.03.005.

Farahany, N.A., 2023. The battle for your brain: defending the right to think freely in the age of neurotechnology. St. Martin’s Press.

Gallistel, C.R., King, A.P., 2011. Memory and the computational brain: Why cognitive science will transform neuroscience. John Wiley Sons.

Gladstone, B., Neufeld, J., 2011. The Influencing Machine. W.W. Norton.

Hartmann, T., Wirth, W., Schramm, H., Klimmt, C., Vorderer, P., Gysbers, A., Sacau, A.M., 2016. The spatial presence experience scale (spes): Measuring spatial presence, engagement, and immersion in virtual environments. Journal of Media Psychology 28, 1–15. doi:10.1027/1864-1105/a000137.

Hartmann, W.R., Klapper, D., 2018. Super bowl ads. Marketing Science 37, 78–96. doi:10.1287/mksc.2017.1055.

Herman, E.S., Chomsky, N., 1988. Manufacturing Consent: The Political Economy of the Mass Media. Pantheon Books.

JASP Team, 2024. Jasp (version 0.19.0) [computer software].

Jeon, M., Lim, S., Lapinski, M.K., Bente, G., Spates, S.A., Schmaezle, R., 2024. Attention and retention effects of culturally targeted billboard messages: An eye-tracking study using immersive virtual reality. bioRxiv.

Kim, H.K., Park, J., Choi, Y., Choe, M., 2018. Virtual reality sickness questionnaire (vrsq): Motion sickness measurement index in a virtual reality environment. Applied Ergonomics 69, 66–73. doi:10.1016/j.apergo.2017.12.016.

Kingstone, A., Smilek, D., Ristic, J., Kelland Friesen, C., Eastwood, J.D., 2003. Attention, researchers! it is time to take a look at the real world. Current Directions in Psychological Science 12, 176–180. doi:10.1111/1467-8721.01255.

Kramer, A.D., Guillory, J.E., Hancock, J.T., 2014. Experimental evidence of massive-scale emotional contagion through social networks. Proceedings of the National Academy of Sciences 111, 8788–8790. doi:10.1073/pnas.1320040111.

Kümmerer, M., Bethge, M., 2023. Predicting visual fixations. Annual Review of Vision Science 9, 269–291.

Lasswell, H.D., 1927. Propaganda technique in the World War. Knopf.

Lim, S., Cho, H.J., Jeon, M., Cui, X., Schmaelzle, R., 2024. Using vr and eyetracking to study attention to and retention of ai-generated ads in outdoor advertising environments. bioRxiv doi:10.1101/2024.08.15.607684.

Lleras, A., Buetti, S., Xu, Z.J., 2022. Incorporating the properties of peripheral vision into theories of visual search. Nature Reviews Psychology 1, 590–604. doi:10.1038/s44159-022-00097-1.

Matz, S.C., Kosinski, M., Nave, G., Stillwell, D.J., 2017. Psychological targeting as an effective approach to digital mass persuasion. Proceedings of the National Academy of Sciences 114, 12714–12719. doi:10.1073/pnas.1710966114.

McCombs, M.E., Shaw, D.L., 1972. The agenda-setting function of the mass media. Public Opinion Quarterly 36, 176–187.

McGuire, W.J., 1968. Personality and attitude change: An information-processing theory. doi:10.1016/b978-1-48323071-9.50013-1.

Miller, L.C., Shaikh, S.J., Jeong, D.C., Wang, L., Gillig, T.K., Godoy, C.G., et al., 2019. Causal inference in generalizable environments: systematic representative design. Psychological Inquiry 30, 173–202. doi:10.1080/1047840X.2019.1693866.

Neijens, P., Araujo, T., Möller, J., de Vreese, C., 2024. Measuring exposure and attention to media and communication: Solutions to wicked problems. Amsterdam University Press.

Neisser, U., 1964. Visual search. Scientific American 210, 94–103.

Nelson, J.L., Webster, J.G., 2016. Audience currencies in the age of big data. International Journal on Media Management 18, 9–24.

Perloff, R., 1999. The third person effect: A critical review and synthesis. Media Psychology 4, 353–378. doi:10.1207/s1532785xmep0104_4.

Potter, J.W., 2008. The importance of considering exposure states when designing survey research studies. Communication Methods and Measures 2, 152–166.

Pärnamets, P., Johansson, P., Hall, L., Balkenius, C., Spivey, M.J., Richardson, D.C., 2015. Biasing moral decisions by exploiting the dynamics of eye gaze. Proceedings of the National Academy of Sciences 112, 4170–4175. doi:10.1073/pnas.1415250112.

Rosenblatt, F., Farrow, J.T., Rhine, S., 1966. The transfer of learned behavior from trained to untrained rats by means of brain extracts. Proceedings of the National Academy of Sciences 55, 548–555.

Schmälzle, R., Huskey, R., 2023. Integrating media content analysis, reception analysis, and media effects studies. Frontiers in Neuroscience 17, 1155750.

Schmälzle, R., Lim, S., Cho, H.J., Wu, J., Bente, G., 2023. Examining the exposure-reception-retention link in realistic communication environments via vr and eye-tracking: The vr billboard paradigm. Plos One 18, e0291924. doi:10.1371/journal.pone.0291924.

Schupp, H., Cuthbert, B., Bradley, M., Hillman, C., Hamm, A., Lang, P., 2004. Brain processes in emotional perception: Motivated attention. Cognition and Emotion 18, 593–611.

Sherman, B.E., Turk-Browne, N.B., 2024. 587 Attention and Memory. Oxford University Press.

Skinner, B.F., 1958. Teaching machines. Science 128, 969–977.

Skurka, C., Keating, D.M., 2024. How repeated exposure to persuasive messaging shapes message responses over time: a longitudinal experiment. Human Communication Research, hqae008.

Slater, M.D., 2004. Operationalizing and analyzing exposure: The foundation of media effects research. Journalism Mass Communication Quarterly 81, 168–183. doi:10.1177/107769900408100112.

Sutherland, M., 2020. Advertising and the mind of the consumer: what works, what doesn’t and why. Routledge. doi:10.4324/9781003114833.

Tausk, V., 1919. Über die entstehung des beeinflussungsapparates” in der schizophrenie. Internationale Zeitschrift für Psychoanalyse 5, 1–33.

Todd, R.M., Manaligod, M.G., 2018. Implicit guidance of attention: The priority state space framework. Cortex 102, 121–138.

Turow, J., 2017. The Aisles Have Eyes: How Retailers Track Your Shopping, Strip Your Privacy, and Define Your Power. Yale University Press.

Turow, J., 2022. Media Today: Mass Communication in a Converging World. Routledge. doi:10.4324/9781003133933.

Valkenburg, P.M., Peter, J., Walther, J.B., 2016. Media effects: Theory and research. Annual Review of Psychology 67, 315–338.

Zajonc, R.B., 1968. Attitudinal effects of mere exposure. Journal of Personality and Social Psychology 9, 1. doi:10.1037/h0025848.

Zillmann, D., Bryant, J., 1985. Affect, mood, and emotion as determinants of selective exposure, in: Zillmann, D., Bryant, J. (Eds.), Selective Exposure to Communication. Lawrence Erlbaum Associates, Hillsdale, NJ, pp. 157–190.

